# Deep Learning Driven Cell-Type-Specific Embedding for Inference of Single-Cell Co-expression Networks

**DOI:** 10.1101/2024.08.12.607542

**Authors:** Yuting Bai, Kun Qian, Qiang Lin, Wei Fan, Rui Qin, Bing He, Feng Ding, Wanfei Liu, Peng Cui

**Author notes:** Equal contribution (Yuting Bai, Kun Qian, Qiang Lin).

## Abstract

The inference of gene co-expression module in specific cell types is important for understanding of cell-type-specific biological processes. We have developed DeepCSCN, an unsupervised deep-learning framework, to infer gene co-expression modules from single-cell RNA sequencing (scRNA-seq) data. Utilizing a global-to-local network construction approach, DeepCSCN can infer co-expression at the whole sample level and construct cell-type-specific co-expression networks. Systematic evaluations on eight public scRNA-seq datasets show that DeepCSCN significantly outperforms eight existing methods in co-expression network construction. Furthermore, DeepCSCN effectively identifies cell-type-specific co-expression networks that are more enriched for cell-specific functional pathways compared to current methods. Finally, application of DeepCSCN on public scRNA-seq data revealed 280 cell-specific gene modules across 27 cell types, including epithelial cells, immune cells, and myonuclei, demonstrating its versatility and accuracy in elucidating cell-type-specific gene co-expression regulation. DeepCSCN offers a powerful tool for researchers to dissect the intricate gene co-expression networks within distinct cell types.

## Introduction

Gene co-expression analysis is crucial for unraveling complex biological processes^1^. Traditionally, microarray or bulk RNA-seq data have been the primary sources for such analyses, showing excellent results in module detection^2^, disease discovery^1, 3^, and identifying genes linked to various biological processes^4, 5^. Recent advancements in single-cell RNA sequencing (scRNA-seq) technology now allow for gene expression analysis at single-cell resolution, which uncovers heterogeneity within cell populations. However, the high throughput, noise, and sparsity associated with scRNA-seq data complicate network construction, making it difficult to accurately infer gene co-expression from this type of data.

Multiple methods utilizing various correlation metrics have been developed to infer co-expression relationships from scRNA-seq data, including Pearson^3^, Spearman^3^, GENIE3^6^, PIDC^7^, minet^8^, glasso^9^, and sclink^10^. For instance, the popular Weighted Gene Co-expression Network Analysis (WGCNA)^3^ employs the Pearson coefficient as a co-expression correlation metric and primarily used for constructing gene co-expression networks from bulk RNA-seq data. PIDC and minet utilize mutual information (MI) to measure co-expression. GENIE3 uses a tree-based integration approach to estimate gene correlations and has recently been applied to single-cell expression data^11^. Glasso estimates a sparse inverse covariance matrix from Gaussian data, and its variants have been used to tackle different challenges in gene network construction^12, 13^. Sclink refines the Gaussian graphical model for gene co-expression estimation by explicitly modeling scRNA-seq data.

However, due to the curse of dimensionality and high sparsity of scRNA-seq data, these methods exhibit low accuracy in co-expression construction. To solve the issue, some pre-processing methods were used to correct the raw scRNA-seq data. For instance, pseudo-bulk reduces data sparsity by averaging cell expressions through clustering. However, it often yields worse estimations compared to raw data^14^. Additionally, aggregation decreases the number of samples, resulting in the loss of statistical information and negatively impacting analysis. Imputation is used to correct scRNA-seq data by filling in missing values. However, these methods can introduce biased estimates of gene expression and may lead to false-positive gene correlations^15, 16^. Therefore, these correction methods do not significantly improve the performance of single-cell co-expression estimation. These methods also struggle with large-scale transcriptome integration data, which involves multiple sources and high heterogeneity, making it difficult to identify gene modules with high correlation.

Additionally, current methods struggle to accurately infer cell-type-specific gene networks for large datasets. Recently published methods, sclink and CS-CORE^17^, directly screen highly expressed genes of specific cell types for the network construction. However, this approach often overlooks some cell-type-specific lowly-expressed genes, leading to inaccuracies in final constructed networks. Therefore, there is an urgent need to develop new computational methods that can effectively extract and integrate co-expression information from large scRNA-seq datasets to improve the accuracy of co-expression prediction. Furthermore, we need to enhance the current methods to infer more precise cell-type-specific co-expression networks and reveal their underlying gene co-expression patterns.

Here, we developed an unsupervised deep-learning framework, DeepCSCN, to construct gene co-expression networks from scRNA-seq data. DeepCSCN accurately infers cell-type-specific co-expression networks from large samples by employing features decoupling of cell types. Our evaluations show that DeepCSCN outperforms existing methods. We successfully applied DeepCSCN to construct cell-type-specific networks across 27 cell types including epithelial cells, immune cells, and myonuclei. DeepCSCN offers several innovations and advantages. (1) Effective Feature Extraction: DeepCSCN utilizes unsupervised learning for gene feature extraction, addressing the sparsity issue of scRNA-seq data and improving the accuracy of gene correlation calculations. (2) Higher Accuracy: DeepCSCN achieves higher accuracy in gene co-expression prediction and identifies more cell-type-specific gene modules compared to the existing methods. (3) Global-to-Local Network Construction: DeepCSCN first trains on all samples to extract gene embeddings, then selects cell-type-specific dimensions from these embeddings based on feature disentanglement. This approach enables the inference of co-expression networks from a whole-sample level to a specific cell type level. In summary, our work introduces a novel method for dissecting the intricate gene co-expression networks within distinct cell types, providing insights into cell-type-specific biological processes.

## Results

### Overview of DeepCSCN

We developed a deep leaning framework, named DeepCSCN, to construct cell-type-specific gene co-expression networks from scRNA-seq data (Fig.1). To leverage the consistency between co-expression and clustering analyses in identifying gene groups with similar expression patterns, we transformed the gene co-expression module identification task into an unsupervised clustering task within DeepCSCN. To achieve this, we first developed the deep clustering model GeneCluster to learn a robust gene feature representations from scRNA-seq data (Methods). Given an scRNA-seq matrix *x* ∈ *R*^*n*\**m*^, where n represents the number of input genes and m represents the number of cells, GeneCluster learns a mapping function *f*_*θ*_(*x*_*n*_) that can generate low-dimensional gene embeddings to predict latent gene-gene correlations. GeneCluster integrates feature extraction with clustering to extract cluster-guided feature embeddings. Using these embeddings, we computed gene correlations using cosine similarity to construct co-expression networks. Next, based on the extraction of cell-type-specific embeddings, we constructed cell-type-specific gene co-expression networks. We validated DeepCSCN using multiple scRNA-seq datasets, demonstrating that it significantly improves the accuracy of co-expression predictions, compared to existing methods. Finally, we applied DeepCSCN to build large-scale cell-type-specific gene co-expression network atlases using human scRNA-seq data from 27 different cell types.

**Fig 1.**
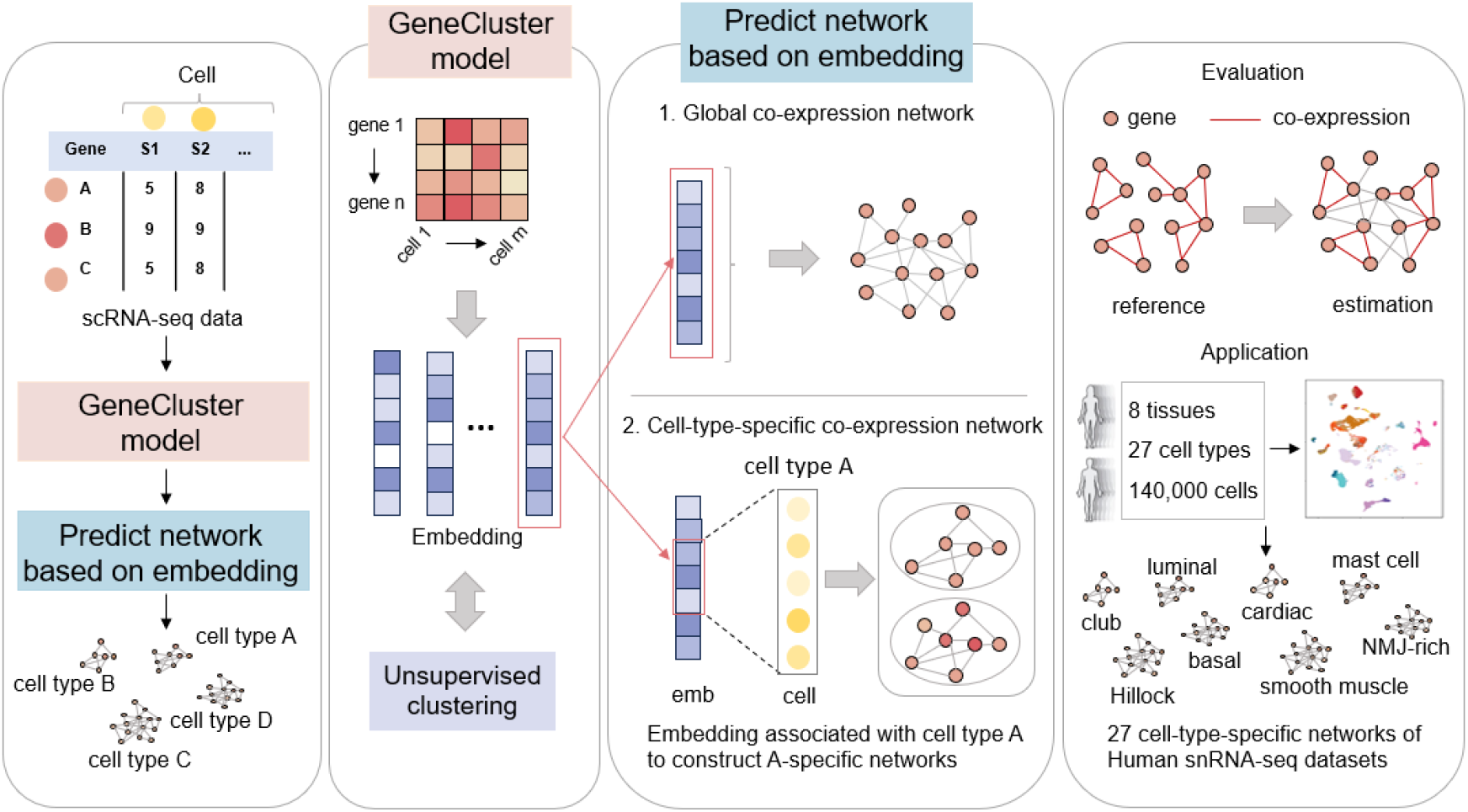
The framework and evaluation of DeepCSCN model. Using gene expression matrix from scRNA-seq data as input, the GeneCluster model was trained to learn gene embedding representations, which were then used to predict co-expression networks. The process involves two main steps. 1 Global Co-expression Network Prediction: Initially, embeddings from all samples are used to predict the global co-expression networks. 2 Cell-type-Specific Network Construction: In the second step, cell-type-specific embeddings are extracted to construct gene co-expression networks tailored to each cell type. For example, embeddings specific to cell type A are utilized to construct co-expression networks for cell type A. The known co-expression relationships were used for evaluation. Finally, the DeepCSCN model was applied to build 27 cell-type-specific networks from human scRNA-seq datasets.

### Evaluation of GeneCluster’s clustering performance

Since GeneCluster is essentially a clustering model, it is necessary to evaluate its performance of both feature extraction and clustering. To achieve this, we used visualization and computational clustering assessment metrics to compare GeneCluster with four commonly used gene expression clustering methods, including K-Means^18^, Hclust^19^, Fanny^20^, and Mclust^21^.

We visualized the clustering results of different models using Uniform Manifold Approximation and Projection (UMAP)^22^ on eight human and mouse scRNA-seq datasets (Supplementary Table 1). For GeneCluster, the gene representations learned from model training were used to visualize. In contrast, for the other four clustering methods, the original expression data were directly used to visualize. To determine the optimal number of clusters for the gene representations obtained by GeneCluster, we used the Bayesian Information Criterion (BIC) and applied the same number of clusters across all methods. The results indicated that K-Means, Hclust, Fanny, and Mclust showed weak separation between clusters, while GeneCluster demonstrated clear and distinct clusters (Fig.2a). This trend was consistently observed across all eight datasets (Supplementary Fig.1), suggesting that GeneCluster extracts more effective gene representations, thereby enhancing clustering performance.

**Fig 2.**
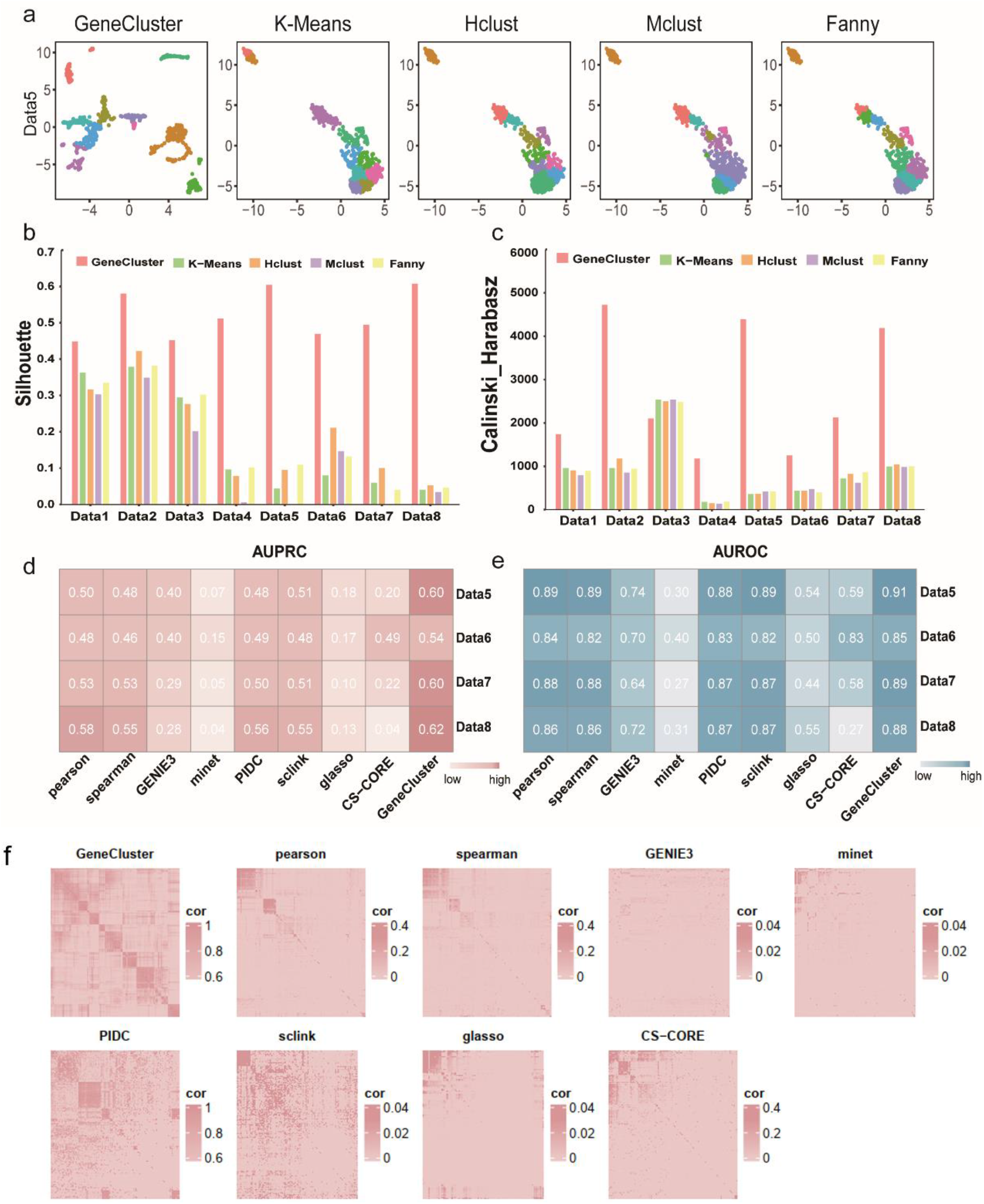
Evaluation of clustering and gene co-expression performances. **a**. UMAP visualization of Data5. **b**. Comparison of Silhouette among different clustering methods. **c**. Comparison of Calinski-Harabasz among different clustering methods. **d**. AUPRC comparison between DeepCSCN and the eight other methods. **e**. AUROC comparison between DeepCSCN and the eight other methods. **f**. Comparison of correlation heatmaps for different co-expression methods (Rows and columns are clustered by default).

Moreover, by calculating the Silhouette coefficient^23^ and Calinski-Harabasz (CH)^24^ index as clustering metrics (Methods), GeneCluster demonstrated significantly higher values for both metrics compared to other clustering methods (Fig.2b-c). These results highlight that GeneCluster excels in feature extraction and clustering, suggesting its effectiveness in integrating these processes into a unified framework. Unlike traditional methods that separate representation learning from clustering, GeneCluster simultaneously learns data representation and performs clustering. This integrated approach leads to the extraction of cluster-oriented gene features, resulting in substantial improvements in gene clustering.

### Evaluation of gene co-expression metrics

After extracting gene expression features using GeneCluster, we aimed to construct co-expression matrices based on these features. To determine the most suitable metric for co-expression, we evaluated five commonly used correlation metrics: cosine similarity, covariance, Euclidean distance, Pearson correlation and Spearman correlation. We selected four human PBMC scRNA-seq datasets for testing, as these datasets contain validated co-expression relationships and were also used in a recent benchmark study comparing co-expression methods^14^. Our evaluation showed that cosine similarity exhibited the highest co-expression accuracy (Supplementary Fig.3) when tested against known co-expression relationships. Consequently, we chose cosine similarity for the co-expression assessment for our calculation.

Next, we constructed the co-expression matrices for 400 highly variable genes from each dataset and compared them with eight other co-expression metrics or methods: Pearson, Spearman, GENIE3, minet, PIDC, sclink, glasso, and CS-CORE. We used two evaluation metrics: the precision-recall curve (AUPRC) and the area under the receiver operating characteristic curve (AUROC), to assess the performance. GeneCluster achieved the highest AUPRC scores (Fig.2d) and AUROC scores (Fig. 2e) across the four datasets compared to the other methods, indicating higher accuracy in calculating gene co-expression. Notably, glasso and minet had the lowest co-expression scores, suggesting that the penalized Gaussian graphical model and the pairwise mutual information-based model are less effective for scRNA-seq data. Conversely, PIDC, which uses improved mutual information, performed better. Correlation coefficient methods, such as Pearson and Spearman, also achieved high co-expression scores. Additionally, while all methods showed high AUROC scores, they had lower AUPRC scores. This discrepancy arises from the sparse nature of the ground-truth network, where negative instances (absence of co-expression) vastly outnumber positive instances (presence of co-expression). Since AUROC is sensitive to class imbalance, we also used AUPRC to address this issue by considering both precision and recall values^7^.

Additionally, we visualized the correlation matrices calculated by different methods and found that GeneCluster was able to identify more modules compared to the other methods. In contrast, Pearson, Spearman, and PIDC identified fewer modules, while GENIE3 and glasso failed to identify any modules. These results suggest that traditional co-expression methods, which rely on direct calculations of gene expression counts, are less effective at accurately estimating gene correlations and achieving meaningful clustering. In contrast, GeneCluster leverages deep learning to generate gene representations from scRNA-seq data, rather than relying on direct correlation calculations from expression counts. This approach significantly enhances the accuracy of co-expression calculations.

### Construction of global gene co-expression networks

To demonstrate the application of GeneCluster in constructing global gene co-expression networks, we applied GeneCluster to four human PBMC datasets. After obtaining gene embeddings using GeneCluster and constructing co-expression matrix with cosine similarity, we utilized hierarchical clustering and the dynamic tree cut algorithm^25^ to the matrix to identify gene clustering modules (Methods). In the PBMC1 drop-seq dataset (Data5), we identified 25 highly correlated co-expression modules (Fig.3a), with results for the other three datasets provided in Supplementary Fig. 4.

**Fig 3.**
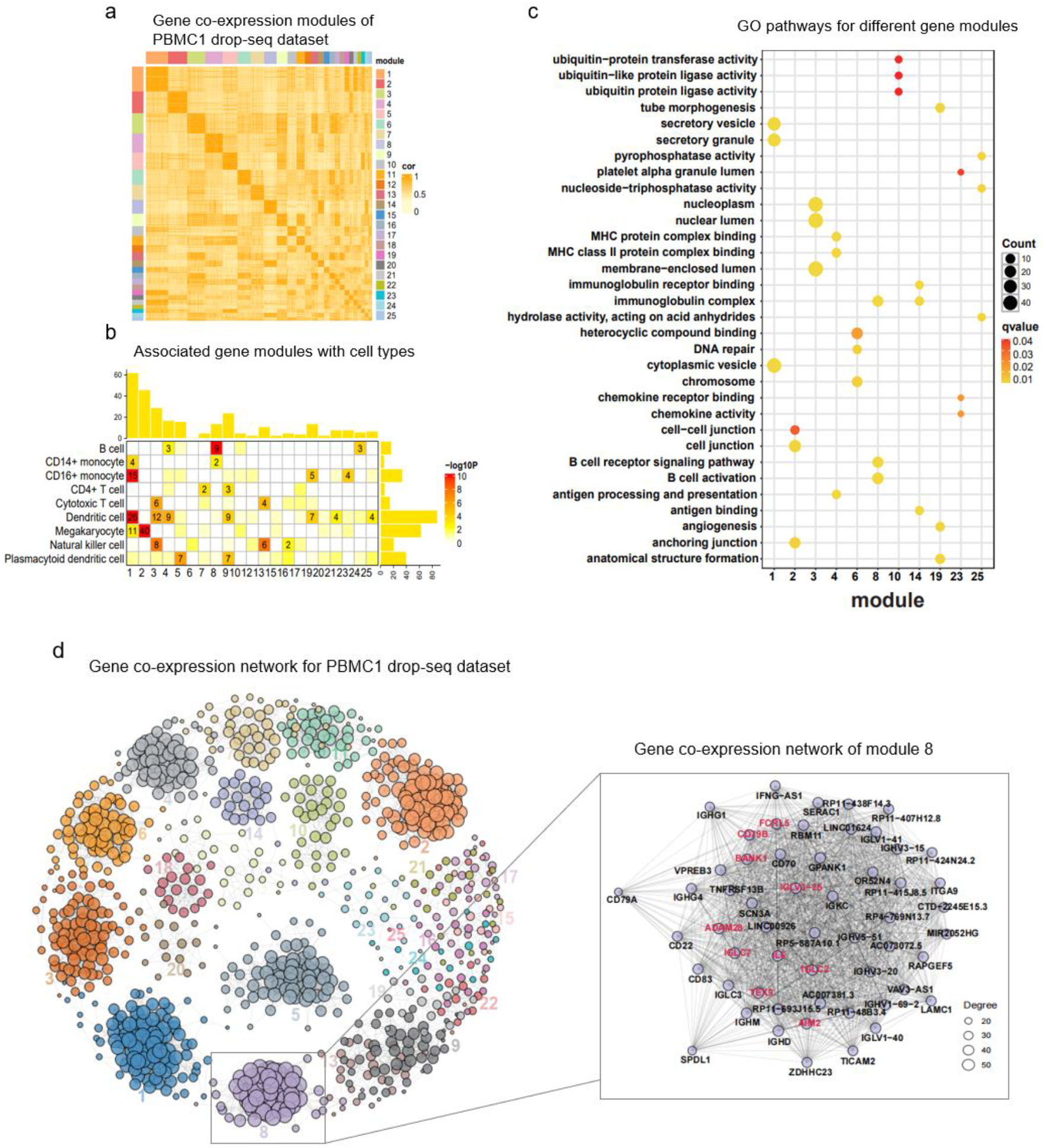
Construction of global gene co-expression networks. **a**. Heatmap of gene co-expression modules for the PBMC1 drop-seq dataset. **b**. Associate gene modules with cell types (the number in the box represents significance, with larger values indicating greater significance). **c**. GO pathways for different modules. **d**. Gene co-expression network for the PBMC1 drop-seq dataset. The rectangular box on the right is a further presentation of the gene co-expression relationship of module 8, with the hub gene highlighted in red.

To validate the accuracy of co-expression module identification, we first examined whether these modules are associated with specific types of cells. We used the FindMarkers function in Seurat^26^ to identify marker genes for each cell type. Then, we conducted a hypergeometric test to analyze the association between these markers and gene module groups. Our analysis revealed that many gene modules were significantly associated with specific cell types (p-value < 0.05) (Fig.3b). For examples, module 1 was significantly enriched in CD16+ monocytes and Dendritic cells, module 2 in Megakaryocytes, and Module 8 in B cell. Further gene ontology (GO) enrichment analysis^27^ of the different gene modules revealed significant enrichment in multiple functional pathways related to cell-type-specific functions (Fig.3c). Detailed analysis of module 8 indicated significant enrichment in pathways associated with B cell activation and the B cell receptor signaling pathway. The gene relationships within module 8 are shown in Fig.3d, with *FCRL5, IGLC2, CD79B, IGLC7*, and *IL6* identified as hub genes due to their high degree of co-expression. *CD79B* are known to be expressed exclusively in B cells and B cell malignancies, and antibodies against these proteins are useful markers for diagnosing precursor B acute lymphoblastic leukemia^28, 29^. FCRL5 is a protein expressed in specific types of memory B cells following exposure to certain infections or pathogens, making it a promising target in vaccine development^30^. AIM2 is highly expressed in B cells of lupus patients and promotes B cell differentiation by modulating the Bcl-6-Blimp-1 axis, potentially representing a novel therapeutic target for systemic lupus erythematosus^31^. By constructing a global gene co-expression network and associating the modules with cell types and specific molecular functions, we demonstrated the capability of GeneCluster to produce accurate gene co-expression networks.

### CONSTRUCTION of cell-type-specific gene co-expression networks

Although global gene modules can be associated with specific cell types, they do not precisely assign each module to a specific cell type. To address this, we developed a novel method for constructing cell-type-specific gene modules. First, we generated an association matrix to quantify the importance of gene embeddings in distinguishing different cell types (Fig.4a). This was achieved by calculating the correlation between the expression matrix of cell-type-specific genes (i.e. differentially expressed genes among different cell types) and the gene embedding matrix obtained from GeneCluster training. Using the association matrix, we then extracted the top 100-dimensional gene embeddings for each cell type and calculated co-expression to construct cell-type-specific networks based on these extracted embeddings.

Using our newly developed algorithm, we constructed cell-type-specific networks from the human PBMC1 drop-seq dataset. We focused on five cell types, including B cell, CD4+ T cell, CD16+ monocyte, Cytotoxic T cell and Natural killer cell. We first calculated the gene correlation matrix for the top 100-dimensional features of each cell type separately. For B cell, the matrix was clustered into two groups, with group 2 being more significantly cell-type-specific score (see in Methods) and notably enriched with most B cell marker genes (Fig.4b). Consequently, genes within group 2 were thought to be B cell-specific. We then calculated the correlation matrix for genes in group 2 and identified the modules using dynamic tree cutting, resulting in a total of four B cell-related modules (Fig.4c). Each module was enriched for B cell-associated pathways. For example, module 1 was primarily associated with MHC protein complex binding, module 2 was mainly linked to B cell activation, and module 3 was enriched in immunoglobulin receptor binding functions (Fig. 4d).

**Fig 4.**
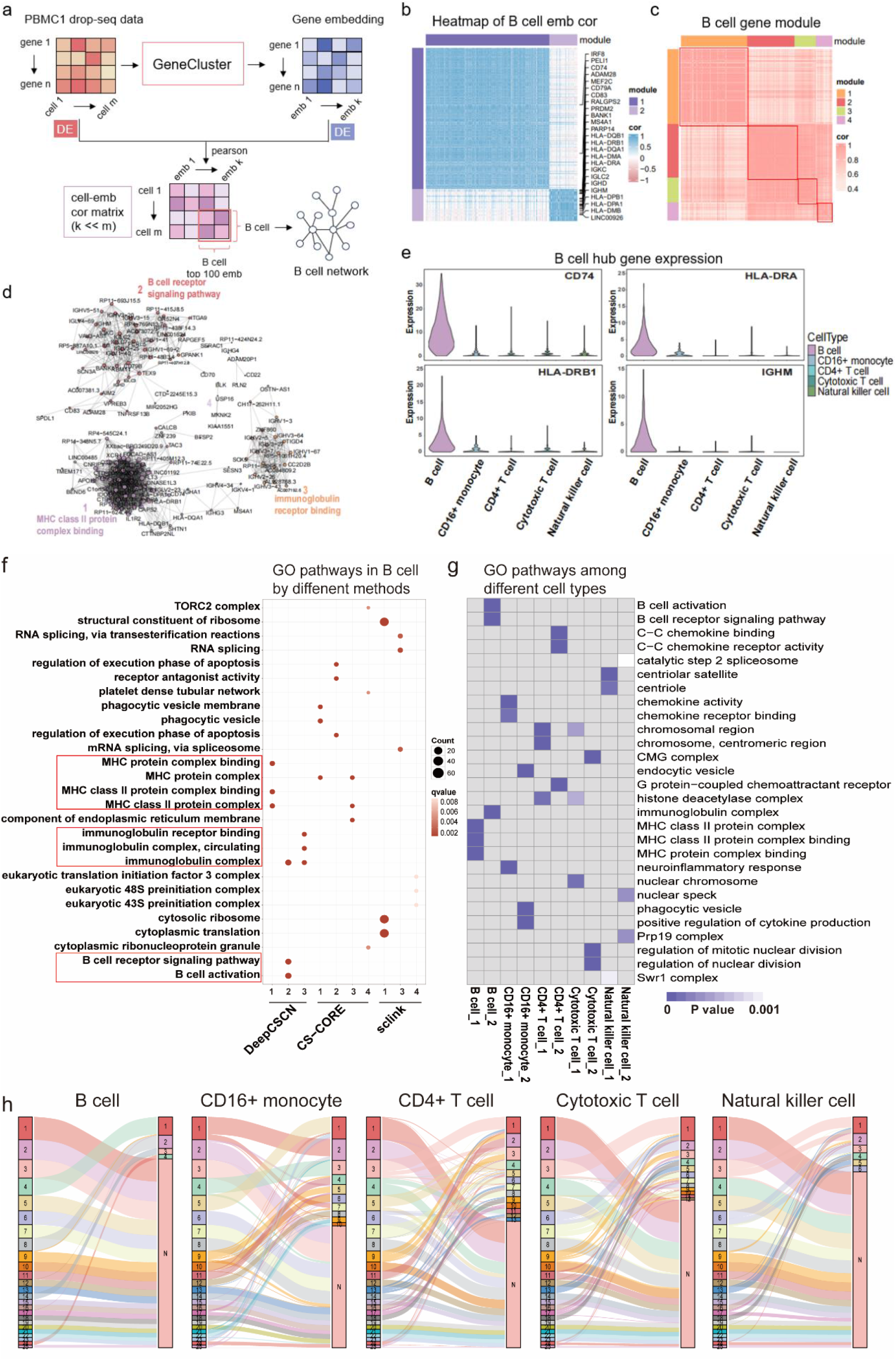
Construction of cell-type-specific gene co-expression networks. **a**. An illustration of cell-type-specific network construction. **b**. B cell top100-dimensional gene embedding correlation heatmap. The genes marked on the right are the B-cell marker genes recognized by the Seurat package. **c**. Co-expression gene modules in B-cell. **d**. Network of B-cell co-expression gene modules. **e**. Expression of B-cell hub genes in different cell types. **f**. Comparison of DeepCSCN, CS-CORE, and sclink for B-cell GO enrichment pathways. **g**. GO pathway analysis of the two largest gene modules for B cell, CD4+ T cell, CD16+ monocyte, Cytotoxic T cell, and Natural killer cell. **h**. The relationship between global co-expression modules and cell-type-specific co-expression modules. The left column is the global module, the right column is the cell-type-specific modules for B cell, CD4+ T cell, CD16+ monocyte, Cytotoxic T cell and Natural killer cell respectively. N denotes gene modules unrelated to specific cell type.

For the other cell types, we identified 13 modules in CD4+ T cell, 10 modules in CD16+ monocyte, 12 modules in Cytotoxic T cell and 6 modules in Natural killer cell (Supplementary Fig.6B). GO enrichment analysis was performed on the specific modules identified for each cell type, and the two largest gene modules are displayed in Fig.4g. We observed strong specificity in the pathways enriched for different cell types. For B cell, both module 1 and module 2 were specifically enriched in B cell pathways. Similarly, the GO term related to “chemokine activity” and “chemokine receptor binding” were specifically enriched in gene module of CD16+ monocyte (Fig.4g). These results further demonstrate the effectiveness of our method in identifying cell-type-specific modules.

For each cell type, we also identified hub genes within each module based on their connectivity in the co-expression network. For example, in B cell, the hub genes for module 1 include *SHTN1, CTTNBP2NL, FLT3, HLA-DRB1, CD74, HLA-DRA, AREG, CAPS2*, and *C1orf54*. The hub genes for module 2 are *VPREB3, AIM2, IGHD, RBM11, TEX9, RP11-48B3*.*4, ADAM28, BANK1*, and *CD79B*. The hub genes for module 3 include *ZNF860, AC007192*.*6, CC2D2B, CD22, IGHV1-3, IGHV1-67, IGHV2-5, IGHV3-43, IGHV3-64*, and *IGHV3-7*. To further validate the specificity of the identified B cell-related modules, we randomly selected four hub genes and assessed their expression levels across different cell types (Fig.4e). We observed that these genes exhibited high expression levels exclusively in B cells, with low or negligible in other cell types. Additionally, most of these hub genes are well-known for their involvement in B cell-specific functional pathways. For example, immunoglobulins (expressed by *IGHV-1-3, IGHV1-67, IGHV2-5*, etc.) play a crucial role in B cell-specific antigen recognition during the recognition phase of humoral immunity^32, 33^.

Furthermore, to validate the effectiveness of our approach in identifying cell-type-specific modules, we compared with two recently published methods: sclink and CS-CORE, which are used for constructing cell-type-specific networks. In constructing of the B-cell-specific network, we ensured a fair comparison by selecting 1,000 genes for each method based on their respective gene screening criteria (Methods). Our method identified four B cell-related modules, with three of these modules being enriched in B cell-related pathways. In contrast, CS-CORE also identified four modules, but only module 1 and 3 were enriched in MHC protein complex pathways related to B cells. Sclink identified 60 modules, none of which were enriched in B cell-related pathways (Fig.4f, Supplementary Table 3). Therefore, our method outperforms existing approaches in constructing cell-type-specific networks.

Additionally, we compared the differences between the global gene co-expression construction and the cell-type-specific construction. We aligned cell-type-specific gene modules onto the global modules. For five cell types, we identified 32 cell-type-specific modules that were not detected by the global network construction. These include 11 CD4+ T cell-specific modules, 7 CD16+ monocyte-specific modules, 10 Cytotoxic T cell-specific modules, 3 Natural killer cell-specific modules and 1 B cell-specific modules (Fig.4h). This result demonstrates that the global gene co-expression construction is not able to precisely identify cell-type-specific gene modules. Therefore, constructing cell-type-specific networks is a more efficient way to cell-type-specific gene modules.

### DeepCSCN identifies gene networks from 27 human cell types

To evaluate the performance of DeepCSCN on large-scale cell atlases and demonstrate its potential application in constructing cell-type-specific gene networks, we applied DeepCSCN to identify gene co-expression modules in a large dataset from the GTEx Portal^34^, which includes human epithelial cells, immune cells, and myonuclei, comprising a total of 137,766 cells and covering 27 cell types (Fig.5a). For each cell type, we selected 1,000 highly variable genes for training, and then extracted gene embeddings for post-training of all samples to construct global co-expression networks. Additionally, we identified the top 100-dimensional gene embeddings associated with specific cell types to construct cell-type-specific networks (Methods) (Fig.5b).

**Fig 5.**
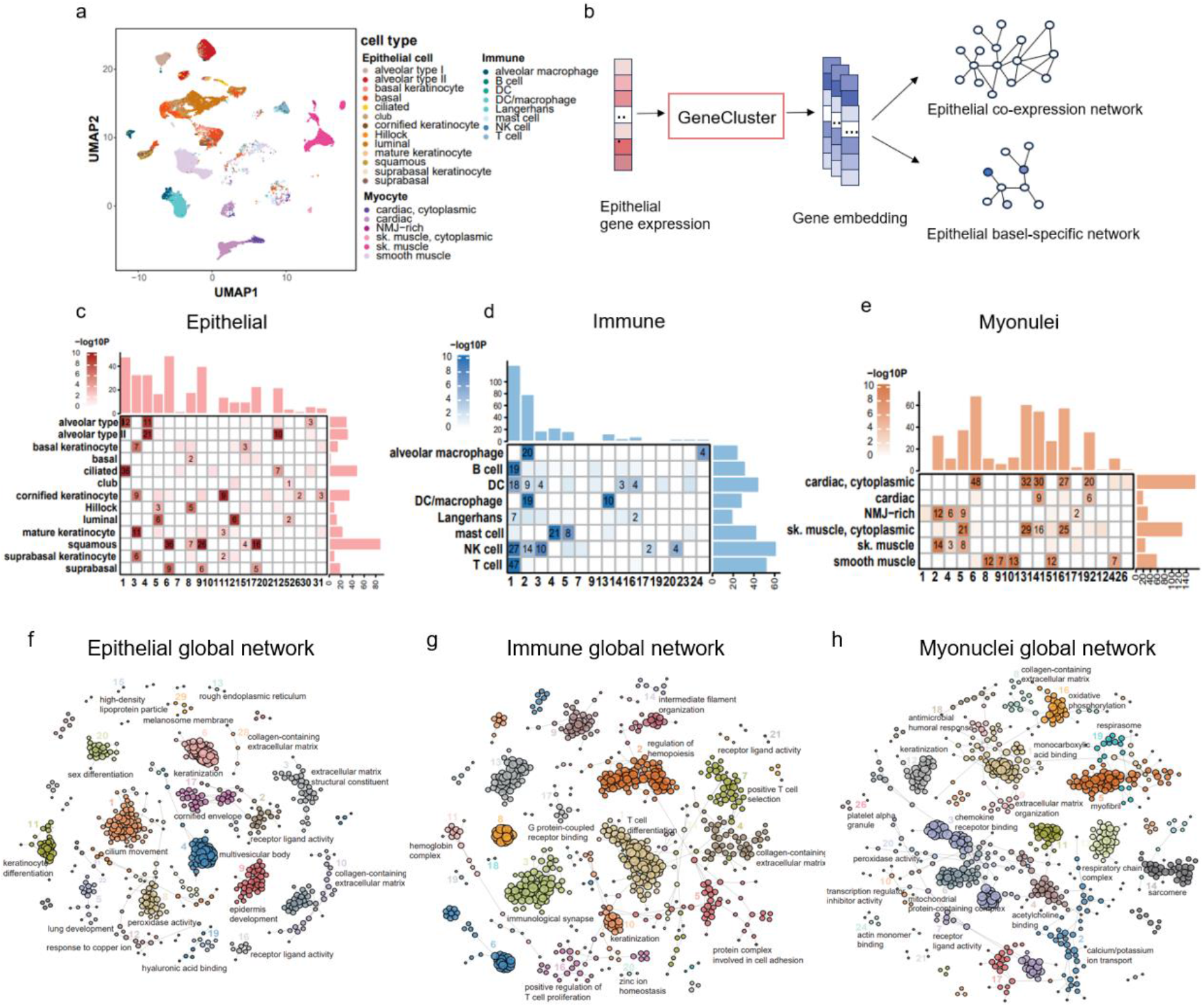
Global gene co-expression networks of the Human scRNA-seq dataset. **a**. UMAP representation of single-nucleus profiles (dots) colored by cell types. **b**. Illustration of DeepCSCN-based human scRNA-seq co-expression network. **c-e**. The global gene co-expression modules of epithelial, immune and myonuclei is associated with cell types (the number in the box represents significance, with larger values indicating greater significance). **f-h**. The co-expression networks of epithelial, immune and myonuclei.

Using DeepCSCN, we first constructed global gene co-expression networks and identified a total of 31, 24, and 26 co-expression modules in epithelial cells, immune cells and myonuclei, respectively. We also used the number of marker genes enriched in the modules to correlate the global gene modules with cell types. In epithelial cells, module 1 is associated with ciliated cells and alveolar type 1 cells, module 3 is associated with mature keratinocytes, cornified keratinocytes, basal keratinocytes and suprabasal keratinocytes, and module 6 is associated with squamous cells and suprabasal cells (Fig.5c). In immune cells, module 1 is associated with T cells, Natural killer (NK) cells, B cells, Dendritic cells (DCs), and Langerhans cells, and module 4 and 5 are associated with mast cells (Fig. 5d). In myonuclei, module 2 and 4 are significantly enriched in NMJ-rich cells and skeletal muscle cells, module 6 is enriched in “cytoplasmic” cardiac cells, and modules 8, 9, 10, 15, and 24 are enriched in smooth muscle cells (Fig.5e). Furthermore, we visualized the complex co-expression network relationships for epithelial cells, immune cells, and myonuclei, highlighting the distinct pathways enriched in each module. For instance, in myonuclei, module 1 is enriched in monocarboxylic acid binding, module 5 is enriched in the myofibril, and module 14 is enriched in the sarcomere (Fig.5f-h). These results demonstrate the strong capability of DeepCSCN in constructing gene co-expression networks from large-scale scRNA-seq datasets.

Subsequently, we conducted cell-type-specific network construction for 13 types of epithelial cells, 8 types of immune cells, and 6 types of myonuclei individually. Initially, we extracted the top 100-dimensional gene embeddings for each cell type, computed their co-expression matrices, performed clustering, and identified modules associated with cell types (Supplementary Fig.7-9). We identified a total of 280 gene modules across 27 different cell types (see Supplementary Table 4), with an average of 10 modules per cell type.

We conducted GO enrichment analysis on these gene modules, selecting the two largest gene modules for each cell type to display. In myonuclei, we found that GO terms related to cardiac muscle tissue development and extracellular matrix structural constituent are only enriched in gene modules of cardiac (Fig.6a). The myonuclei contain six cell types: cardiac cells, “cytoplasmic” cardiac cells, NMJ-rich skeletal muscle cells, skeletal muscle (sk. muscle) cells, “cytoplasmic” skeletal muscle cells, and smooth muscle cells. We identified 11 modules in cardiac cells, 8 modules in “cytoplasmic” cardiac cells, 10 modules in NMJ-rich skeletal muscle cells, 10 modules in skeletal muscle cells, and 18 modules in “cytoplasmic” skeletal muscle cells, respectively (Fig.6b-g). For the epithelial cells (Supplementary Fig.10A), we found that GO terms related to keratinization and keratinocyte differentiation are only enriched in gene module of cornified keratinocyte. The GO term “regulation of glial cell apoptotic process” is only enriched in a gene module of mature keratinocyte, which has a critical role in regulating epidermal development and restraining carcinogenesis^35^. For the immune cells (Supplementary Fig.10B), the enriched GO terms in gene modules are also associated with cell-type-specific biological functions. Detailed enrichment pathways for these modules are provided in Supplementary Table 4.

**Fig 6.**
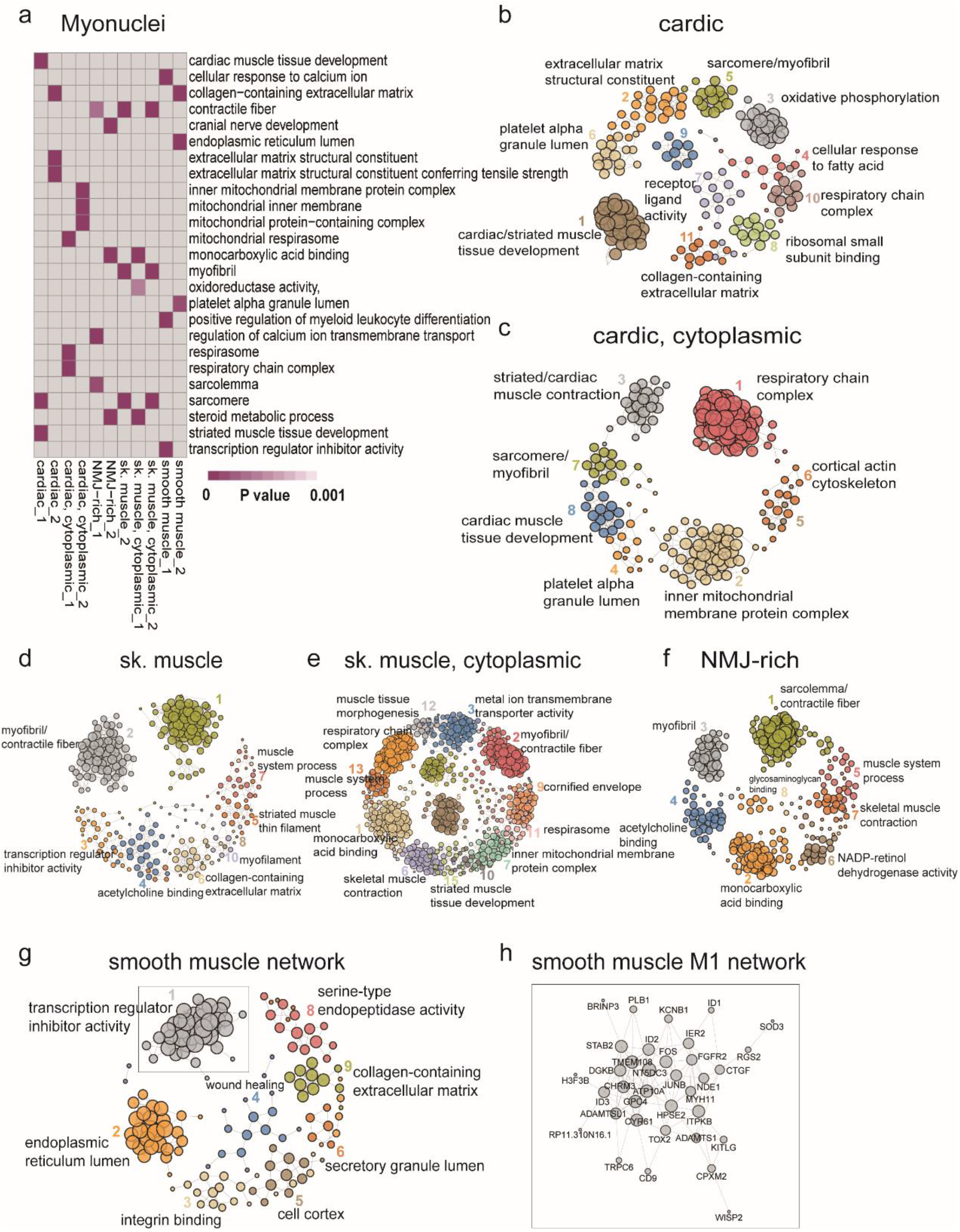
Cell-type-specific gene co-expression networks of the Human scRNA-seq dataset. **a**. Enriched GO terms in the two largest gene modules identified from myonuclei. **b-g**. Network of cardiac cells, “cytoplasmic” cardiac cells, sk. muscle cells, “cytoplasmic” sk. muscle cells, NMJ-rich skeletal muscle cells, and smooth muscle cells. **h**. Network of module 1 in smooth muscle cells.

We described the gene modules for smooth muscle cells in detail as an example. Smooth muscle cells had 9 co-expression modules (Fig.6g). These modules showed the following enrichments: module 1 is mainly enriched to transcription regulator inhibitor activity, module 2 is mainly enriched to endoplasmic reticulum lumen, module 3 is mainly enriched to integrin binding, module 4 is mainly enriched to wound healing, module 5 is mainly enriched in cell cortex, module 6 is mainly enriched in secretory granule lumen, module 8 is mainly enriched in serine-type endopeptidase activity and module 9 is mainly enriched in collagen-containing extracellular matrix. For each module, we identified the top 10 genes with high connectivity as hub genes. In module 1, the hub genes are *ID1, ID2, ID3, ROR2, KITLG, FOS, ITPKB, KCNB1, JUNB*, and *HPSE2* (Fig.6h). Previous research has highlighted their roles in cardiogenesis and vascular system formation, suggesting that *ID* genes are crucial targets during cardiogenesis^36^. By constructing smooth muscle specific networks, we identified key target genes involved in cardiogenesis, which could be significant for future disease treatment.

## Discussion

As scRNA-seq data continue to grow, traditional methods for constructing gene co-expression networks have become inefficient and inaccurate. To address this issue, we developed DeepCSCN, a deep learning framework designed to construct both global co-expression networks and cell-type-specific co-expression networks from scRNA-seq data. At the heart of DeepCSCN is GeneCluster, an unsupervised deep learning model for extracting gene representations from scRNA-seq data. GeneCluster was inspried by DeepCluster^37^ used in image classification area, which optimizes feature extraction through pseudo-labels and clustering tasks. GeneCluster significantly enhances the extraction of gene features, thereby improving the efficiency and accuracy of gene co-expression networks.

Traditional methods for constructing cell-type-specific gene co-expression networks, such as sclink and CS-CORE, typically involve selecting highly expressed genes from each cell type—often the top 1,000 genes. This approach can lead to the inclusion of non-cell-specific genes and the omission of lowly expressed, cell-specific genes, resulting in incomplete and inaccurate networks. In contrast, DeepCSCN integrates gene expression information across different cell types to derive comprehensive cell-type-specific gene features. By disentangling gene representations learned by GeneCluster, and calculating correlations between individual cell gene expression values and dimensions of gene embeddings, we accurately identified gene features specific to each cell type. This approach allows for more precise and comprehensive construction of cell-type-specific gene networks compared to existing methods.

In summary, our study introduces the DeepCSCN framework for constructing cell-type-specific gene co-expression networks. Compared to other methods, DeepCSCN demonstrates superior performance in clustering accuracy and network construction. We successfully applied DeepCSCN to construct 27 cell-type-specific networks across nearly 140,000 cells, highlighting its potential for large-scale scRNA-seq atlas analysis. We believe DeepCSCN offers a robust and effective tool for inferring cell-type-specific co-expression networks, thereby enhancing our ability to characterize biological pathways and molecular mechanisms at the cell-type level.

## Methods

### Overview of DeepCSCN (Cell-type-specific co-expression network inference based on deep learning)

DeepCSCN was designed with three main components: (1) training the GeneCluster model to learn gene embedding representations; (2) constructing a global gene co-expression network based on the representations; (3) disentangling cell-type-specific gene representations and constructing cell-type-specific co-expression networks. Each component is described below.

### Data summary and preprocessing

To evaluate the performance of DeepCSCN, we obtained eight datasets from mouse cortex data^38^ and human peripheral blood mononuclear cells (PBMC) dataset^38^ downloaded from Single Cell Protal. In addition, we also collected human scRNA-seq dataset containing 27 cell types to construct cell-type-specific co-expression networks. The details of these datasets are provided in Supplementary Table 1.

All scRNA-seq datasets were processed using the Scanpy package^39^. For each dataset, genes with expression values of zero in more than 90% of cells were removed, and the top 1000 highly variable genes were selected as input data.

### The GeneCluster framework

GeneCluster includes both a feature extraction module and a clustering module. The feature extraction module utilizes the classical AlexNet^40^ convolutional neural network(CNN) architecture. CNN extract features through convolutional kernels, which provides benefits such as local awareness, parameter sharing, and translation invariance. Given the sparsity of scRNA-seq data, CNN effectively capture local information while filtering out most irrelevant information for single-cell gene feature extraction. To handle datasets with varying sample sizes, pyramid pooling layers^41^ were integrated into the model. The input layer feeds in single-cell gene expression data. In a gene co-expression network, nodes represent genes and edges represent co-expression relationships between genes. Given the scRNA-seq expression data *x*_*n,m*_ ∈ *R*^*N*\**M*^, where N represents the number of input genes, M represents the number of cells. GeneCluster aims to learn a mapping function *f* that can generate low-dimensional gene embeddings E to predict latent gene-gene correlations, i.e.

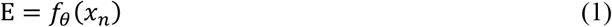

where, X refers to the input vector, specifically the vector of gene expression levels across all cells. (*x*_*n*_) denotes the expression level of the n-th gene, where n ranges from 1 to N, representing the total number of genes, *θ* refers to the learned weights parameters of the function. E denotes the embedding dimension, here equal to 2048 dimensions.

The clustering module functions as a classifier to train the feature extraction module. The output dimension k of the classifier is an adjustable parameter. In each iteration, we used K-means (other clustering methods can also be used) to cluster gene representations derived from the feature extraction module, thereby generating pseudo-labels for the genes. The classifier’s output was then compared with the pseudo-labels from the previous iteration to compute the classification loss, which continuously refines the feature extraction results. The parameters w of the classifier and the parameter *θ* of the mapping are then jointly learned by optimizing the following problem:

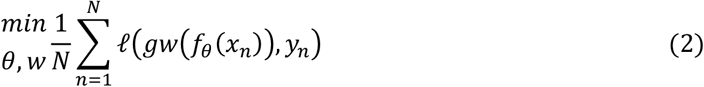

Where ℒ is the cost function. *y*_*n*_ represents the gene’s membership to one of k possible predefined classes. This cost function is minimized using mini-batch stochastic gradient descent and backpropagation to compute the gradient^42^.

In this model, we benefit from using convolutional neural networks to extract sparse single-cell gene features while addressing the limitations of traditional clustering methods, where feature extraction is performed separately from clustering, often leading to ineffective results. GeneCluster integrates the feature extraction model with the clustering model, enabling the extraction of effective gene representations. This integrated approach provides a solid foundation for constructing subsequent cell-type-specific gene co-expression networks.

The GeneCluster model includes two loss functions: clustering loss and classification loss. The clustering loss aims to cluster similar samples together during the training process, thereby improving clustering performance. This loss function is the same as the one used in K-Means clustering. Classification loss is employed to train the model and optimize gene representations. In this study, the Focal loss^43^ function is employed to address the issue of class imbalance.

#### (1) Clustering loss

The clustering loss used in this model is the loss function of K-Means. *f*_*θ*_(*x*_*n*_) is the low-dimensional vector obtained by CNN feature extraction, C is the clustering center matrix; d is the dimension of the gene, i.e., the size of sample; k is the number of cluster centers, and *y*_*n*_ denotes the cluster to which the sample n belongs to, and it is the k-dimensional column vector composed of 0s and 1s (if the sample n belongs to the 0th cluster, *y*_*n*_ = [1,0,0,0 0]T).

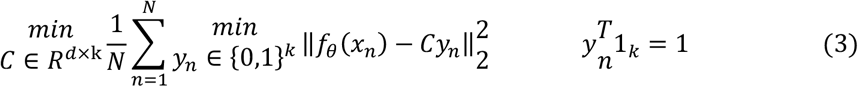

#### (2) Focal loss

Where a_t_ denotes the category weight of the t^th^ sample, y is the true label, 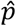 is the predicted probability magnitude, and p_t_ reflects the proximity to the true label y. The larger *p*_*t*_ indicates that it is closer to the category y, i.e., the more accurate the classification.

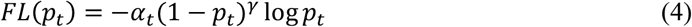

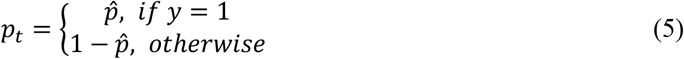

### Construction of global gene co-expression networks

To apply GeneCluster to co-expression analysis, three main steps were carried out. First, low-dimensional feature embeddings of genes were extracted to pre-train the GeneCluster model. Next, using these embeddings, the cosine similarity between genes was computed to generate the correlation matrix. Finally, this correlation matrix was subjected to clustering to derive the co-expression modules.

### Construction of cell-type-specific co-expression networks

The construction of cell-type-specific networks involves three main steps: (1) Extraction of cell-type-specific embeddings; (2) Screening of cell-type-related modules; and (3) Construction of cell-type-specific co-expression networks. Each step is described in detail below:

#### (1) Extraction of cell-type-specific embeddings

First, a cell-embedding correlation matrix was created to assess the importance of gene embeddings in differentiating between cell types. This was done by calculating the correlation between the expression matrix of cell-type-specific genes (i.e., differentially expressed genes among various cell types) and the gene embedding matrix derived from GeneCluster training. Based on this correlation matrix, the top 100-dimensional gene embeddings were selected as cell-type-specific embeddings.

#### (2) Screening of cell-type-related modules

Next, cell-type-specific scores were calculated to screen cell-type-related modules. The score is computed in two steps, as details in formulas (6) and (7). Additionally, B-cell marker genes were used to assist in validation, thereby proving the effectiveness of module selection.

**Step 1:** Calculation of gene relative expression levels across different cells.

To determine the relative expression level of a gene in various cells, divide the expression level of the gene in each cell by the sum of the gene’s expression levels across all cells. This step normalizes the gene expression data, allowing for comparisons between different cellular contexts. The formula for the relative expression level (*E*_*rg*_) of gene *g* in cell *c* is given by:

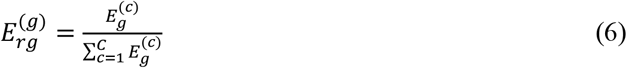

where 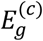 is the expression level of gene *g* in cell *c*, and *C* is the total number of cells.

**Step 2:** Calculation of cell-type-specific score across modules.

To ascertain the cell-type-specific score of genes within a module in a particular cell type, sum the expression levels of the genes in the module for the specified cell type and then divide by the number of genes in that module. This provides a measure of the collective expression activity of the module in different cell types. The formula for the cell-type-specific score (*S*^(*t,m*)^) in cell type *t* for module *m* is:

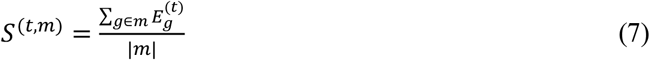

where 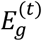 is the expression level of gene *g* in cell type *t*, and ∣*m*∣ denotes the number of genes of module *m*.

#### (3) Construction of cell-type-specific networks

Finally, hierarchical clustering and dynamic shear tree were used to identify cell-type-specific co-expression modules.

### Co-expression analysis of scRNA-seq atlas data

To perform co-expression analysis on the human scRNA-seq atlas, 1,000 highly variable genes were selected from epithelial cells, immune cells, and myonuclei. These genes were input into the GeneCluster model for training. Based on the resulting gene embeddings, we constructed global gene co-expression networks for each cell type: epithelial cells, immune cells, and myonuclei. We then calculated the cell-embedding correlation matrix to evaluate feature importance for each cell type. The top 100-dimensional embeddings for each cell type were defined as cell-type-specific features. Using these features, we computed correlations and performed hierarchical clustering, followed by the identification of cell-type-specific gene modules using formulas (6) and (7). In total, we constructed 27 cell-type-specific co-expression networks covering epithelial cells, immune cells, and myonuclei.

### Evaluation metrics

#### Clustering evaluation

The choice of clustering method can significantly impact the results and significance of the analysis, so it should be carefully considered. To evaluate the effectiveness of the GeneCluster clustering model, we used the Uniform Manifold Approximation and Projection (UMAP) dimensionality reduction method to visualize different clustering approaches. The GeneCluster model employed the extracted 2048-dimensional features and applied UMAP for nonlinear dimensionality reduction, presenting the results in a two-dimensional space. For comparison, other clustering methods, including K-Means, Hclust, Fanny, and Mclust, were also visualized using UMAP-based two-dimensional projections of the expression data to assess their effectiveness.

#### Silhouette Coefficient score

Let *a* is the mean distance between a sample and all other points in the same class, and *b* is the mean distance between a sample and all other points in the next nearest cluster. The Silhouette Coefficient *s* for a single sample is then given as:

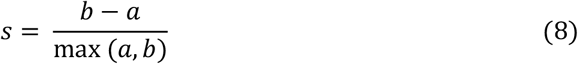

and the Silhouette Coefficient score for a set of samples is given as the mean of each sample’s Silhouette Coefficient score.

#### Calinski-Harabasz score

For a dataset of size *n* which has *k* cell types, the Calinski-Harabas score *s* is defined as the ratio of the between-clusters dispersion mean and the within-cluster dispersion:

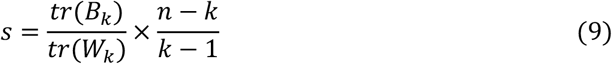

where tr(*B*_*k*_) is trace of the between group dispersion matrix and tr(*W*_*k*_) is the trace of the within cluster dispersion matrix.

### Evaluation of gene co-expression

To validate the effectiveness of co-expression analysis using the DeepCSCN model, eight different co-expression methods were evaluated. AUROC and AUPRC curves are the most commonly used metrics for gene co-expression assessment. The number of correctly (and incorrectly) assigned edges were identified by setting different thresholds and comparing the inferred co-expression relationships with the true network. AUROC plots the false positive rate (FPR) against the true positive rate (TPR), while AUPRC represents the area under the curve defined by recall and precision rates.

### Benchmarking gene co-expression methods

DeepCSCN was compared with eight alternative methods on the experimental data to evaluate their accuracy in constructing gene co-expression networks, including Pearson^3^, Spearman^3^, GENIE3^6^, PIDC^7^, minet^8^, glasso^9^, sclink^10^ and CS-CORE^17^. The Pearson and Spearman were evaluated with the R function “cor”. The method GENIE3 was computed with the R package “GENIE3” (v.1.18.0). The method PIDC was computed with the R package “PIDC” (v.0.3.9). The minet was calculated using the R package “minet” (v.3.54.0). The glasso was calculated using the R package “QUIC” (v.1.1.1). Sclink was implemented from https://github.com/Vivianstats/scLink/. CS-CORE was implementation from https://changsubiostats.github.io/CS-CORE/.

To assess the performance of these co-expression methods, the known gene-gene interaction networks focusing on co-expression related to known transcription factors (TFs) were used as ground truth networks. Specifically, for mouse datasets, the DNA-binding TF co-expression network presented by Zhou^44^ was used as the reference. For human datasets, the gene co-expression discovered in ChIP-seq experiments by Rouillard^45^ was used.

### Computational resources

Model training and all analyses presented in the study were performed on a workstation with an Intel Xeon Gold 6226R CPU with 64 2.9-GHz physical cores addressing 128 GB RAM and an NVIDIA GeForce RTX 3090 GPU addressing 24 GB RAM.

## Data availability

All data used in this work are publicly available, including mouse cortex data, human peripheral blood mononuclear cells (PBMC) data and human scRNA-seq dataset. The datasets were mainly download from GTEx Portal (www.gtexportal.org) and Single Cell Portal (https://singlecell.broadinstitute.org/single_cell). Further details on these datasets are provided in Supplementary Table 1. For functional enrichment analysis, Gene Ontology Database provided used in our analysis is provided by R package clusterProfiler(v.4.2.2)^27^.

## Code availability

DeepCSCN is available on GitHub (https://github.com/Byting820/DeepCSCN). Complete documentation is provided, including model parameters, cluster evaluation, co-expression evaluation, and cell-type-specific co-expression network construction.

## Acknowledgements

This work was supported by Shenzhen Overseas High-Level Talent (Peacock Plan) Project (Grant No. KQTD20180411143628272), STI 2030 – Major Projects and Shenzhen Science and Technology Program (LCYX20220620105200001).

## Author contributions

P.C., W.L., F.D. and Q.L. conceived this project. Y.B. and K.Q. designed and developed the algorithm and model. Y.B. collected and preprocessed the data. Y.B. and K.Q. trained the model.

Y.B. and Q.L. did method evaluation. W.F., R.Q., B.H., W.L. and P.C. provided advice on framework design and model evaluation. Y.B., K.Q., Q.L., F.D., W.L. and P.C. wrote and revised the manuscripts. All authors read and approved the final draft of the paper.

## Additional information

Correspondence and requests for materials should be addressed to P.C..

