## Supplemental Figures for "Deep Learning Driven Cell-Type-Specific Embedding for Inference of Single-Cell Co-expression Networks"

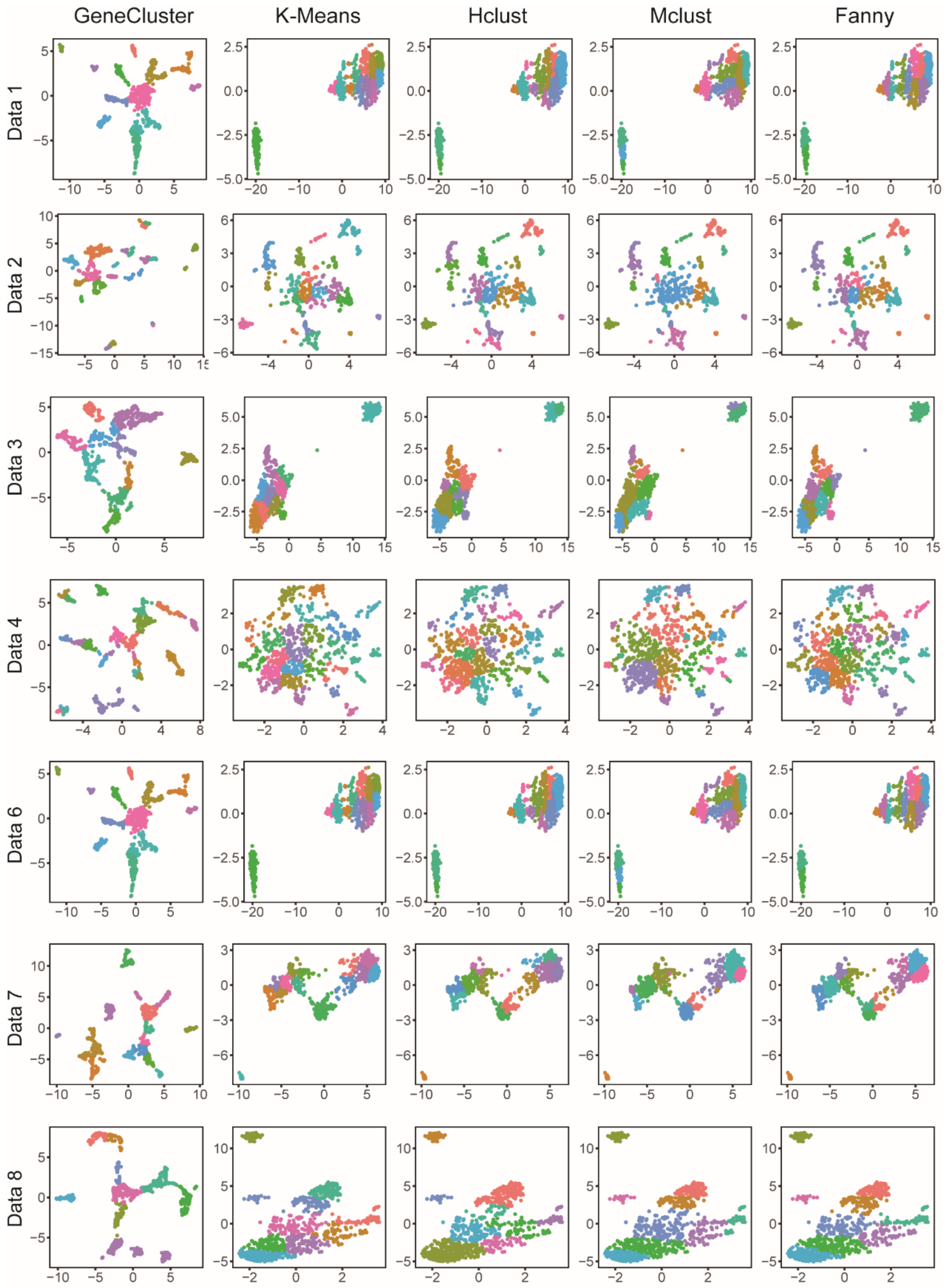


**Supplementary Figure 1.** UMAP Visualization of different clustering models of Data1, Data2, Data3, Data4, Data6, Data7, Data8. Different colors represent different clusters.

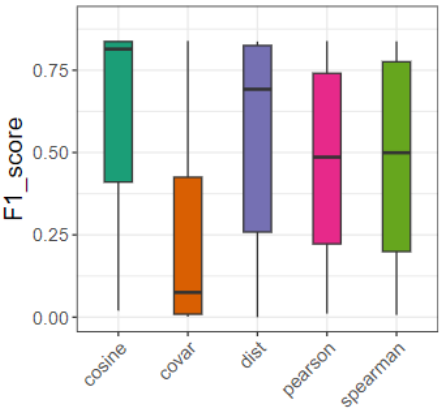


**Supplementary Figure 2.** F1 scores across different correlation metrics.

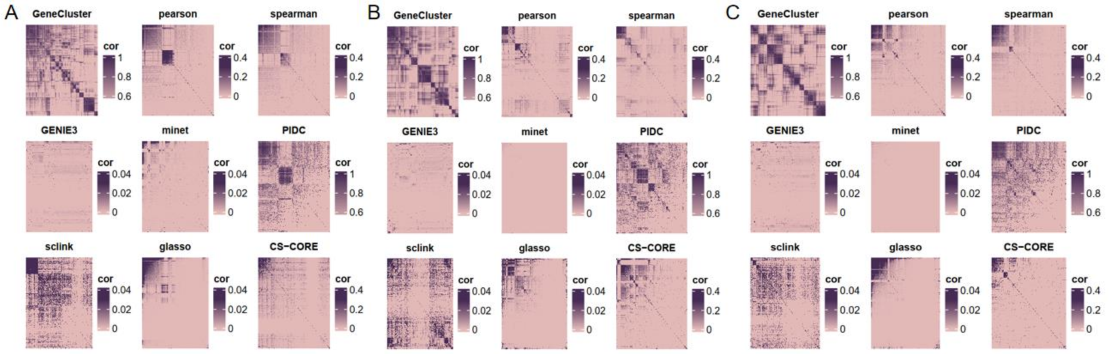


**Supplementary Figure 3.** Comparison of correlation heatmaps for different co-expression methods in (A)PBMC1 inDrops datasets, (B)PBMC2 Drop dataset and (C)PBMC2 inDrop dataset. (Rows and columns are clustered by default).

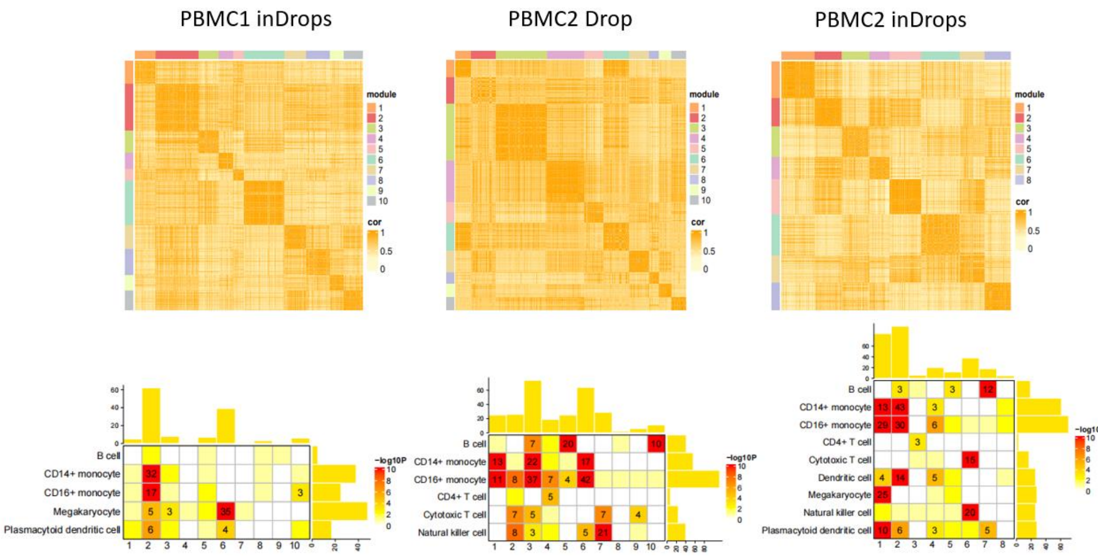


**Supplementary Figure 4.** Heatmap of gene co-expression modules and associate gene modules with cell types for the PBMC1 inDrops, PBMC2 drop-seq and PBMC2 inDrops dataset.

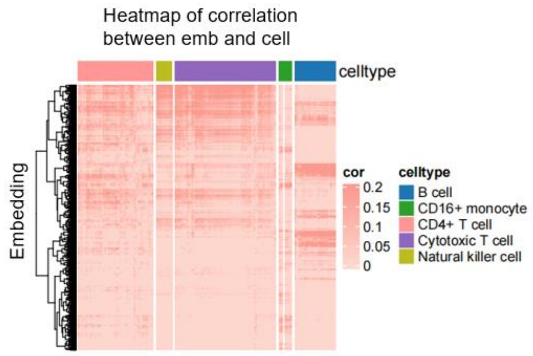


**Supplementary Figure 5.** Heatmap of correlation between gene embedding and cell types.

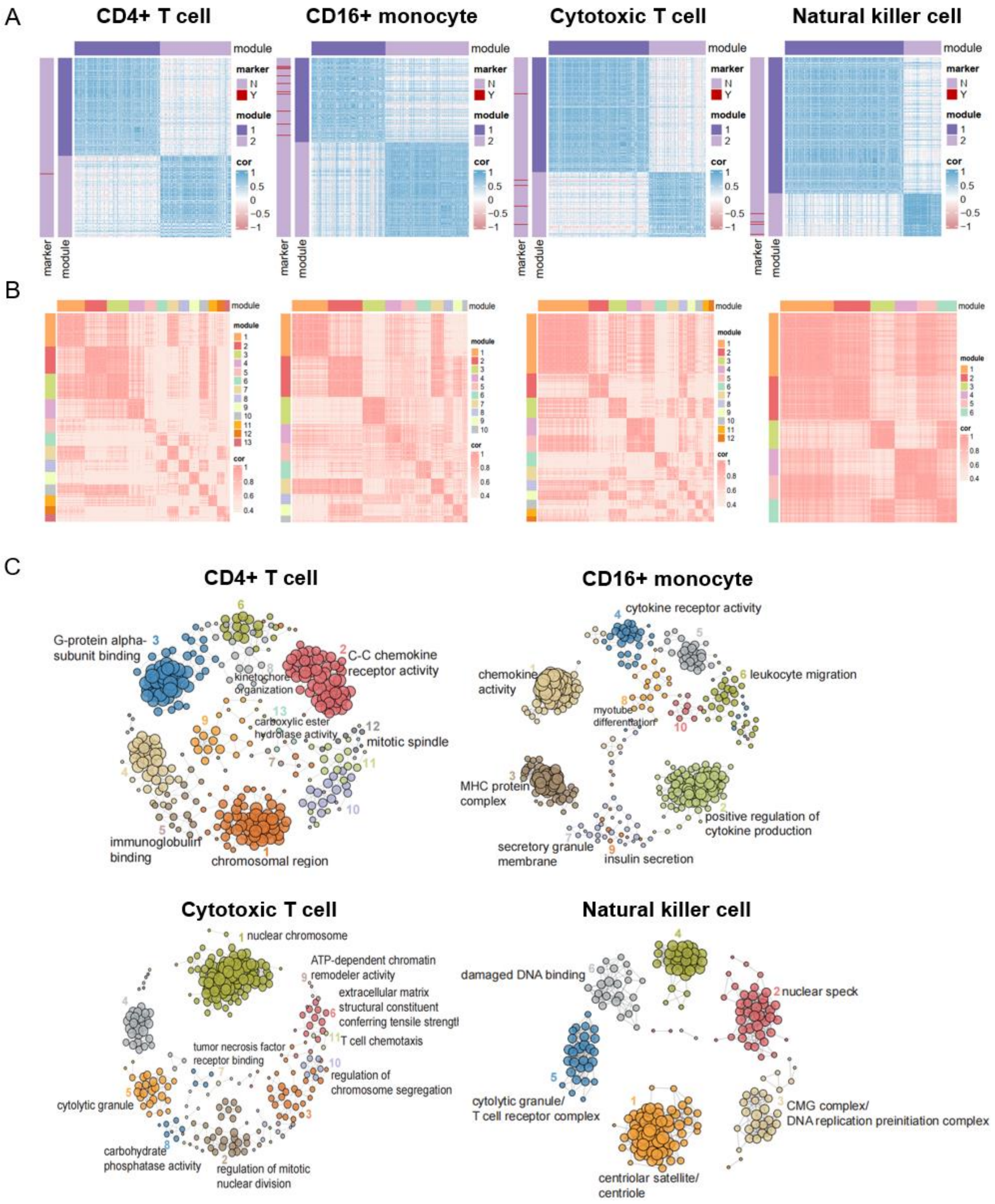


**Supplementary Figure 6. A.** The top100-dimensional feature correlation heatmaps of CD4+ T cell, CD16+ monocyte, Cytotoxic T cell and Natural killer cell. **B.** Co-expression gene modules in CD4+ T cell, CD16+ monocyte, Cytotoxic T cell and Natural killer cell. **C.** The relationship between global co-expression gene modules and cell-type-specific gene modules. D. Network of CD4+ T cell, CD16+ monocyte, Cytotoxic T cell and Natural killer cell co-expression gene modules.

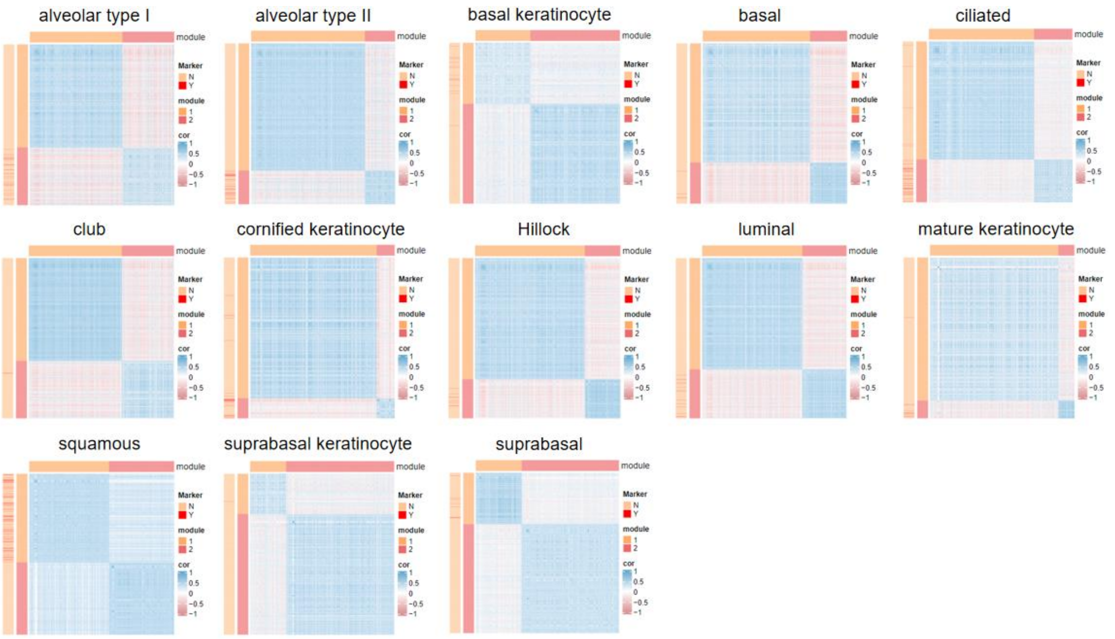


**Supplementary Figure 7.** Gene correlation heatmap visualization of top100 feature for different cell types of Epithelial cells. The red lines in the left bar show marker genes enriched to different modules obtained by clustering different cell types.

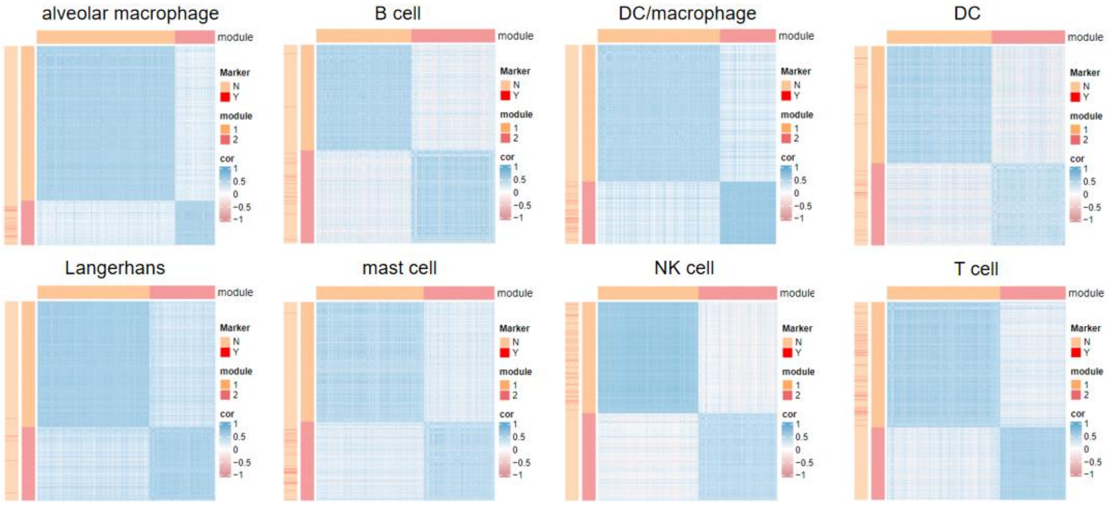


**Supplementary Figure 8.** Gene correlation heatmap visualization of top100 feature for different cell types of Immune cells. The red lines in the left bar show marker genes enriched to different modules obtained by clustering different cell types.

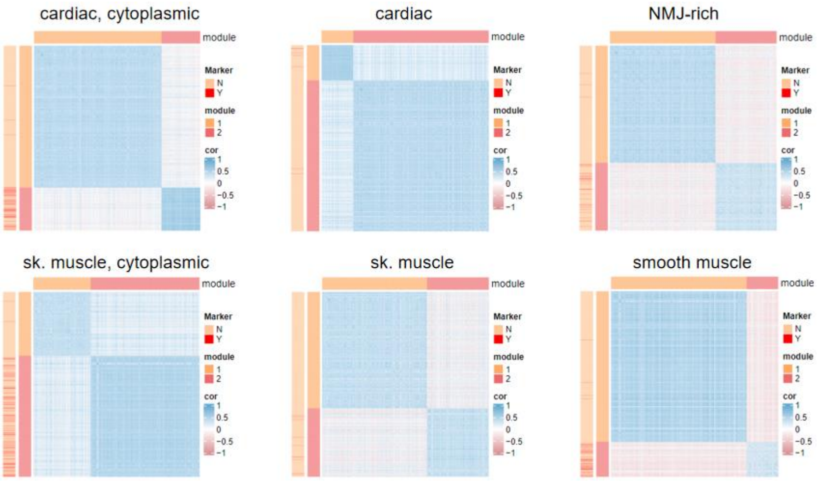


**Supplementary Figure 9.** Gene correlation heatmap visualization of top100 feature for different cell types of Myonuclei. The red lines in the left bar show marker genes enriched to different modules obtained by clustering different cell types.

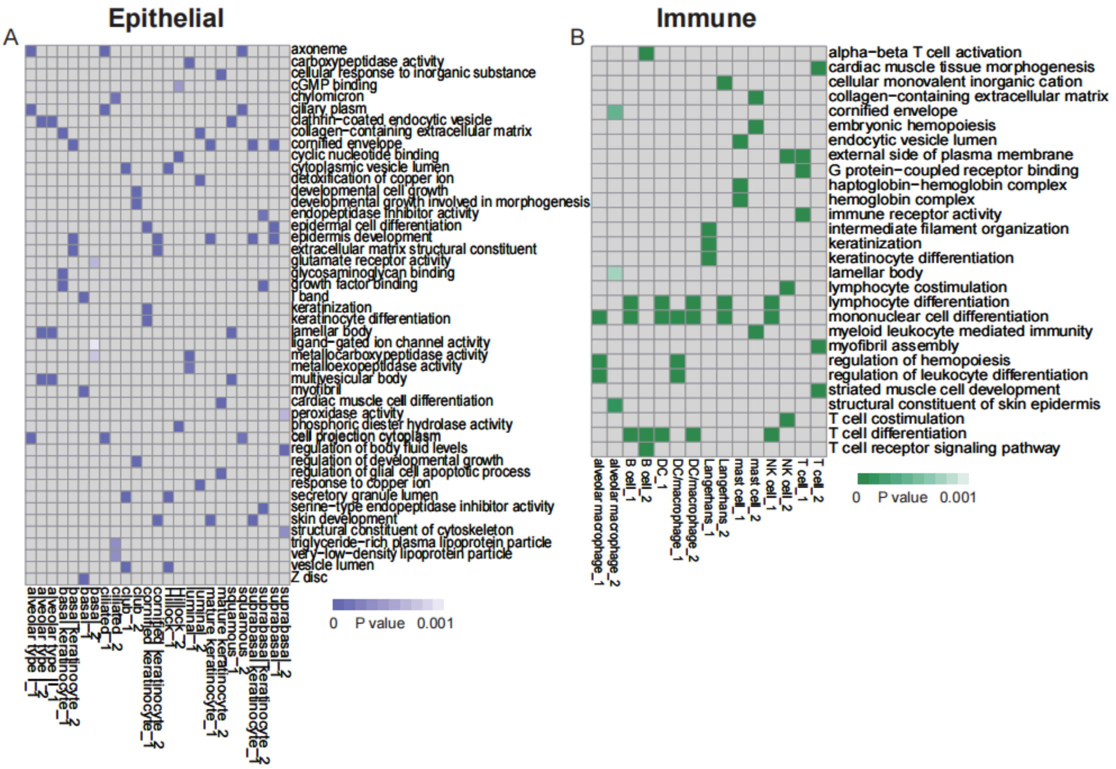


**Supplementary Figure 10.** Enriched GO terms in the two largest gene modules identified from Epithelial and Immune.
